# No Evidence for Systematic White Matter Correlates of Dyslexia: An Activation Likelihood Estimation Meta-Analysis

**DOI:** 10.1101/246009

**Authors:** David Moreau, Josephine E. Stonyer, Nicole S. McKay, Karen E. Waldie

## Abstract

Dyslexia is a prevalent neurodevelopmental disorder, characterized by reading and spelling difficulties. Beyond the behavioral and functional correlates of this condition, a growing number of studies have explored structural differences between individuals with dyslexia and typically developing individuals. To date, findings remain disparate – some studies suggest differences in fractional anisotropy (FA), an indirect measure of white matter integrity, whereas others do not identify significant disparities. Here, we synthesized the existing literature on this topic by conducting a meta-analysis of Diffusion Tensor Imaging (DTI) studies investigating white matter correlates of dyslexia via voxel-based analyses (VBA) of FA. Our results showed no reliable clusters underlying differences between dyslexics and typical individuals, after correcting for multiple comparisons (false discovery rate correction). Because group comparisons might be too coarse to yield subtle differences, we further explored differences in FA as a function of reading ability, measured on a continuous scale. Consistent with our initial findings, reading ability was not associated with reliable differences in white matter integrity. These findings nuance the current view of profound, structural differences underlying reading ability and its associated disorders, and suggest that their neural correlates might be more subtle than previously thought.

## 1. Introduction

Developmental dyslexia is a specific type of learning disability characterized by distinct reading and spelling difficulties. The disorder is typically diagnosed in childhood, affecting around 5-7% of school aged children, and can persist into adulthood (Lindgren, De Renzi, & Richman, 1985; Lyon, Shaywitz, & Shaywitz, 2003; Sally E. Shaywitz, Morris, & Shaywitz, 2008). With heritability estimated to range between 50 and 65% (Habib & Giraud, 2013), dyslexic reading difficulties occur despite appropriate learning environment and adequate resources, and are not attributable to sensory, neurological, psychiatric, intellectual or motivational issues or deficits (Habib & Giraud, 2013; Lyon et al., 2003).

Neuroimaging studies have investigated the neurobiological underpinnings of dyslexia, yielding three key left-hemisphere networks associated with impaired reading. The posterior temporoparietal network has been mainly linked to basic, phoneme level word analysis; the posterior occipitotemporal network, including the visual word form area (VWFA), is commonly associated with word form and fluent reading (Lyon et al., 2003; B. A. Shaywitz et al., 2002; S. E. Shaywitz et al., 1998; Sally E. Shaywitz et al., 2008); whereas the anterior network of the inferior frontal gyrus, including Broca’s area, is involved in speech pronunciation (Sally E. Shaywitz et al., 2008). Furthermore, studies by Shaywitz et al. (1998) and Shaywitz et al. (2002) reported underactivation in posterior temporoparietal and occipitotemporal regions while reading and performing phonological tasks in dyslexics, compared to typical readers. Numerous other functional imaging studies across cultures and stages of development have supported these findings (Brunswick, McCrory, Price, Frith, & Frith, 1999; Horwitz, Rumsey, & Donohue, 1998; Paulesu et al., 2001; Rumsey et al., 1992; Simos, Breier, Fletcher, Bergman, & Papanicolaou, 2000). Shaywitz et al. (1998) and Shaywitz et al. (2002) also observed increased activation of the inferior frontal gyrus, involved in the anterior reading network, among dyslexic compared to typical readers. This hyperactivation is hypothesized to be a compensatory strategy: dyslexic readers use memorization of the structure of words—rather than phonological skills—to read, therefore overengaging frontal brain regions (B. A. Shaywitz et al., 2007; S. E. Shaywitz et al., 2003), though we should note that these findings have been debated in the literature (Hoeft et al., 2007; Norton, Beach, & Gabrieli, 2015; Richlan, 2014; Richlan, Kronbichler, & Wimmer, 2009).

Beyond functional differences, Diffusion Tensor Imaging (DTI) studies have demonstrated impairments in white matter cortical connections between regions among dyslexic readers (Vandermosten, Boets, Wouters, & Ghesquière, 2012). DTI allows probing the distance and direction of water molecule movement, producing form and orientation information about the underlying white matter structures (Assaf & Pasternak, 2008; Soares, Marques, Alves, & Sousa, 2013). In some cortical tissues, such as gray matter and cerebrospinal fluid, diffusion is isotropic; that is, water molecules disperse approximately equally in all directions. Conversely, white matter exhibits anisotropic water movement, with water molecules showing various degrees of diffusion in each direction (Assaf & Pasternak, 2008; Emsell, Van Hecke, & Tournier, 2015; Soares et al., 2013). In typical DTI studies, diffusion images from at least six directions are analyzed using an ellipsoid tensor model—a symmetrical 3x3 matrix. Parallel and perpendicular diffusivities are then calculated and used to estimate properties of underlying tissues. Fractional anisotropy (FA) of the tissue is used most commonly (Assaf & Pasternak, 2008; Soares et al., 2013); FA is measured from 0, isotropic diffusion, to 1, anisotropic diffusion (Assaf & Pasternak, 2008). Other properties include the mean, axial and radial diffusivities (Soares et al., 2013).

Region of interest (ROI) and voxel-based analyses (VBA) can be conducted to compare DTI properties between groups or individuals. In ROI analyses, brain regions defined by *a priori* hypotheses are manually or automatically mapped onto brain images, before the DTI properties of the ROI are averaged within a region and compared across regions. These analyses, however, can be complex, time consuming, and subject to observer and selection biases (Soares et al., 2013; Van Hecke & Emsell, 2015). In contrast, VBA use brain images normalized to a standard brain atlas and smoothed, before computing and comparing DTI properties of each individual voxel. This approach greatly reduces the typical biases of ROI analyses, although this freedom comes at a cost—as VBA is typically less theoretically driven, more drastic corrections for multiple comparisons are often required (Soares et al., 2013; Van Hecke & Emsell, 2015).

Two main avenues of research have been pursued using DTI, employing both ROI and VBA approaches. First, studies have investigated significant differences in FA between dyslexic and typical readers. Two pioneer studies, Klingberg et al. (2000) and Deutsch et al. (2005), identified significant differences in FA in the temporoparietal regions of both hemispheres among small samples of dyslexic and typical reading adults and children, respectively. Lower FA values in the left temporoparietal region among dyslexics compared to typical readers have been further corroborated in subsequent studies (Carter et al., 2009; Rimrodt, Peterson, Denckla, Kaufmann, & Cutting, 2010; Steinbrink et al., 2008), yet despite this apparent convergence, the reported differences within this region vary considerably (Vandermosten et al., 2012). More problematic perhaps, Keller and Just (2009) were unable to replicate these findings in an intervention study, instead reporting lower FA in an anterior region, the left anterior centrum semiovale. Similarly, Koerte et al. (2016) found no significant differences in FA when controlling for false positives adequately. Studies have also found a variety of significant differences in other brain regions, including the superior and inferior frontal regions, precuneus, insula and occipital region in the left hemisphere, superior corona radiata, splenium of the corpus callosum and throughout the right hemisphere (Carter et al., 2009; Deutsch et al., 2005; Frye et al., 2008; Niogi & McCandliss, 2006; Rimrodt et al., 2010; Steinbrink et al., 2008). In addition, Richards et al. (2008) found 45 clusters of significant FA differences between dyslexics and typical readers across the whole brain. Taken together, these findings highlight the wide discrepancies reported in the literature.

Besides group differences contrasting dyslexics with typical readers, additional studies have identified regions where FA values significantly correlate with performance on reading tasks. Numerous studies report positive correlations between FA in the left temporoparietal area of dyslexic or typical readers and reading ability, measured by a range of reading measures (e.g., word reading, pseudo word reading or phonological reading tasks; Beaulieu et al., 2005; Deutsch et al., 2005; Klingberg et al., 2000; Lebel et al., 2013; Nagy, Westerberg, & Klingberg, 2004; Odegard, Farris, Ring, McColl, & Black, 2009; Steinbrink et al., 2008). Similar to the aforementioned literature on group contrasts, however, specific locations within these regions differ considerably between studies (Vandermosten et al., 2012). For example, positive correlations between reading ability and FA have been noted in the superior corona radiata, longitudinal fasciculi, external capsule, centrum semiovale and language areas of the left hemisphere, and bilateral inferior and temporofrontal regions, illustrating the wide variability in results (Deutsch et al., 2005; Keller & Just, 2009; Niogi & McCandliss, 2006; Rimrodt et al., 2010; Steinbrink et al., 2008; Zhang et al., 2014). Finally, negative correlations between reading ability and FA in the posterior/temporal corpus callosum have also been reported (Dougherty et al., 2007; Frye et al., 2008; Odegard et al., 2009; Zhang et al., 2014).

These discrepancies highlight the need to comprehensively examine the variability in brain regions linked to dyslexia. Using Activation Likelihood Estimation (ALE), a technique that determines convergence of activation probabilities across studies (Eickhoff et al., 2009; Eickhoff, Bzdok, Laird, Kurth, & Fox, 2012), Vandermosten et al. (2012) performed a meta-analysis and found that a large cluster (704mm^3^) centered at -29, -17, 26 near the left temporoparietal region, across three DTI studies of correlative and difference that employed VBA. A smaller cluster near the inferior frontal gyrus, centered at -26, 26, 18 was also identified, although less reliably. However, this ALE meta-analysis only examined coordinates where significant differences between dyslexic and typical readers were identified, with particular combinations of studies that created difficulties in interpretation. The software the authors used, GingerALE, has also been updated since, including to correct problems that had a to increase the rate of false positives (Eickhoff, Laird, Fox, Lancaster, & Fox, 2017). Lastly, further correlational research has been conducted since initial publication of the meta-analysis in 2012, suggesting a possible gap in the current literature, and a need to systematically summarize and quantify the relationship between developmental dyslexia and white matter connections. To address these limitations, we conducted a meta-analysis that consisted of two phases. In Phase 1, we focused on differences in FA, assessed via VBA, between dyslexic and typical readers. Phase 2 of the meta-analysis was restricted to correlations between reading ability and VBA studies of FA.

## 2. Results

### 2.1 Study Selection and Characteristics

All details regarding study selection are outlined in Figure 1 (Phase 1) and Figure 2 (Phase 2). Table 1 details the characteristics and demographics of participants included, and the findings of group differences in FA for each study (Phase 1). Table 2 reports the same information for correlations between FA and reading ability (Phase 2).

**Figure 1.**
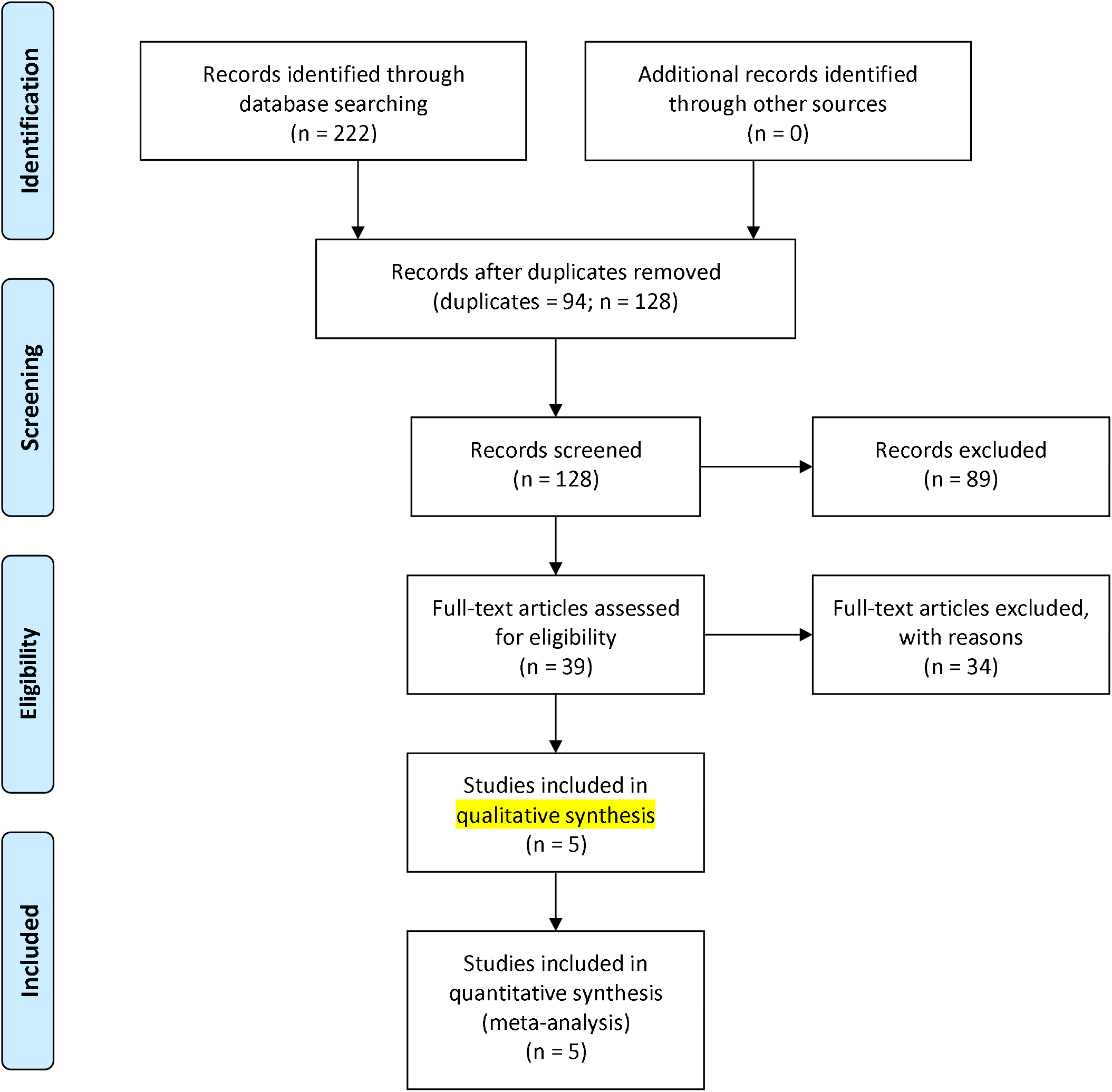
White matter tracts found to have greater fractional anisotropy in controls (A) and in dyslexics (B) across the studies included in Phase 1, uncorrected for multiple comparisons. These tracts are displayed on the skull stripped MNI152 standard brain template at 1mm resolution using FSLview 4.0.1.

**Figure 2.**
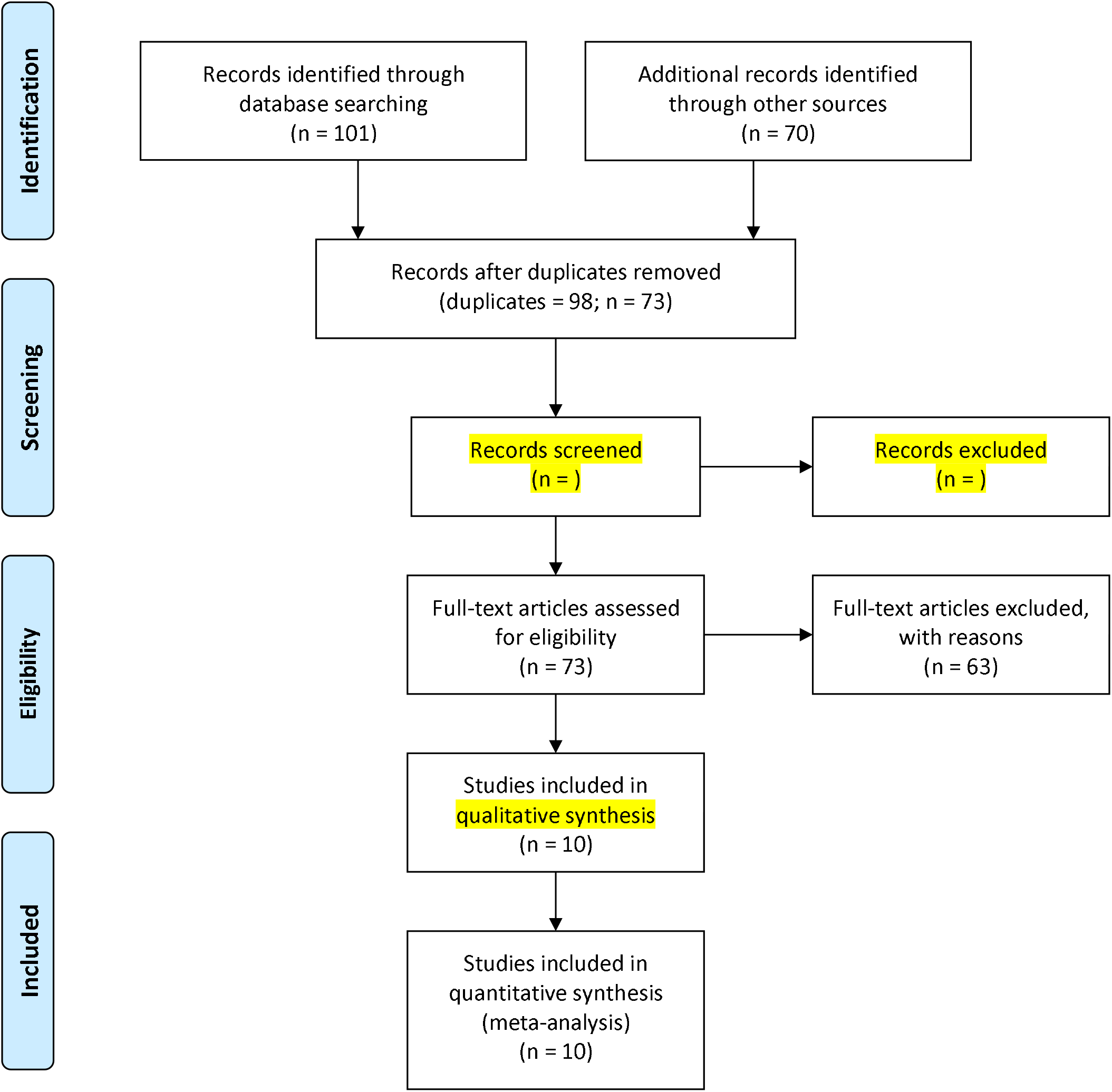
White matter tracts associated with positive (A) and negative (B) correlations between fractional anisotropy and reading ability, uncorrected for multiple comparisons. These tracts are displayed on the skull stripped MNI152 standard brain template at 1mm resolution using FSLview 4.0.1.

### 2.2 Synthesis of Results

Two analyses were run in Phase 1. The analysis of 47 foci from 5 experiments (99 subjects), where FA was significantly greater in typical compared to dyslexic readers, yielded no significant clusters when using a FDR correction of .05. Similarly, the analysis of 17 foci from 2 experiments (52 subjects), where FA was significantly greater in dyslexic compared to typical readers, produced no significant clusters when using a FDR correction of .05.

In Phase 2, two analyses were also undertaken. The analysis of 42 foci from 9 experiments (500 subjects), where reading ability was significantly positively correlated with FA generated no significant clusters when using a FDR correction of .05. Similarly, the analysis of 2 foci from 2 experiments (40 subjects) where reading ability was significantly negatively correlated with FA found no significant clusters when using a FDR correction of .05.

All analyses (Phases 1 and 2) were also run without correcting for multiple comparisons using an uncorrected *p*-value threshold of .05. Tables 3 and 4 present the coordinates of the clusters and their respective contributors for Phase 1 and Phase 2, respectively. Figures 3 and 4 present the white matter tracts corresponding to the uncorrected clusters, for each phase. These include clusters extracted from contrasting dyslexics and control individuals (Phase 1), and those extracted from correlating FA and reading ability (Phase 2). Probabilistic values were computed for each cluster, to determine the most likely white matter tracts associated with each cluster. For details about the clusters, foci and probabilistic estimates, see the Supplemental Material and online analyses outputs.

**Figure 3.**
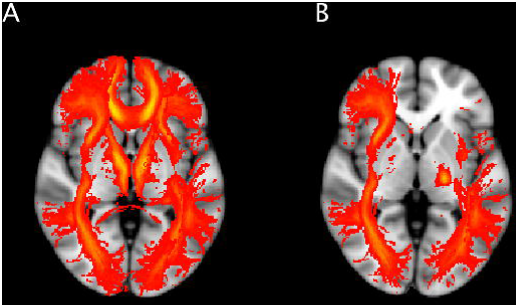
Phase 1 flow diagram.

**Figure 4.**
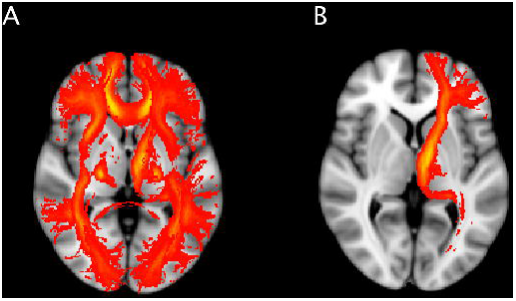
Phase 2 flow diagram.

### 2.3 Additional Analysis

One additional analysis was conducted in Phase 1 where studies including adults and children were analyzed separately. No significant clusters were produced when typical readers had significantly greater FA than dyslexic readers or when dyslexic readers had significantly greater FA than typical readers among studies of adults or children, using an FDR correction of .05.

In Phase 2, several additional analyses were undertaken. Firstly, studies including adult and child participants were again analyzed separately. No significant clusters were produced when reading ability was negatively correlated with FA among adult or child studies, or when reading ability was positively correlated with FA among adult studies, using an FDR correction of .05. One significant cluster—256mm^3^, from MNI coordinates (-22, -10, 32) to (-16, -6, 38), centered at (-18.9, -8.34, 34.9)—was found when analyzing children studies reporting coordinates of positive correlations between reading and FA, but no significant cluster was found when results from Zhang et al. (2014) were excluded from analyses.

## 3. Discussion

We conducted a meta-analysis with two distinct but complementary phases—the first one sought to identify differences in FA between dyslexic and typical readers, whereas the aim of the second phase was to explore correlations between FA and reading ability. Our results showed no systematic differences in fractional anisotropy between dyslexic and typical readers, or as a function of reading ability, after correcting for multiple comparisons. As we argue henceforth, this finding may reflect the current lack of reliable and consistent structural differences in white matter tracts associated with reading ability, or reading disability.

We initially selected and analyzed DTI studies that used VBA to identify cortical coordinates where significant differences in FA existed between dyslexic and typical readers (Phase 1). All studies identified coordinates where typical readers showed greater FA than dyslexic readers, and two studies also reported coordinates where dyslexic readers had greater FA than typical readers. These coordinates were analyzed separately; in both analyses, no reliable clusters were found after correcting for multiple comparisons. That is, no reliable differences in white matter integrity, as measured by FA, were found between dyslexic and typical readers.

Because group comparisons may be too coarse to detect subtle differences, and given that several studies also analyzed the relationship between FA and reading ability on a continuum, we then focused on correlations (Phase 2). Specifically, we selected DTI studies that used VBA to locate cortical coordinates where FA significantly correlated with reading ability or performance on a reading-based task. From this search, nine studies reported coordinates of significant positive FA-reading correlations and two studies reported significant negative correlations. Consistent with Phase 1 findings, no reliable clusters were detected in positive or negative correlation analyses after correcting for multiple comparisons. In other words, reading ability was not reliably associated with white matter integrity as measured by FA.

Our findings nuance the current view of profound differences in FA underlying reading ability and associated disorders such as dyslexia, and correlative relationships between reading ability and FA. We found no reliable differences in FA between dyslexic and typical readers, a finding that contrasts with early findings by Klingberg et al. (2000) and Deutsch et al. (2005) of a temporoparietal difference in FA between dyslexic and typical readers (see also Carter et al., 2009; Rimrodt et al., 2010; Steinbrink et al., 2008). Similarly, the null findings in Phase 2 of this meta-analysis do not support the idea of a relationship between reading ability and FA in the left temporoparietal region, and negative correlation between reading ability and FA in the corpus callosum (Beaulieu et al., 2005; Deutsch et al., 2005; Dougherty et al., 2007; Frye et al., 2008; Klingberg et al., 2000; Nagy et al., 2004).

In line with our findings, however, Vandermosten et al. (2012) noted that despite appearing consistent, each one of the studies they included in their meta-analysis produced coordinates at different locations within the temporoparietal region and corpus callosum; for example, some studies reported correlations with the temporal corpus callosum and others with the posterior corpus callosum. In fact, many studies have also reported differences and correlations in a range of other regions distributed widely throughout the cortex (Deutsch et al., 2005; Frye et al., 2008; Niogi & McCandliss, 2006; Richards et al., 2008; Rimrodt et al., 2010; Steinbrink et al., 2008). Our null results reflect these within-region differences—when aggregated, the inconsistencies across published findings led to disparate clusters that could have emerged stochastically, given the number of comparisons performed in these analyses. We should also point out that the way we conducted the present meta-analysis, looking at aggregates but also at subsets via group splits (e.g., analyzing separately positive and negative correlations in Phase 2) is bound to inflate the false positive rate. This was deliberate on our part, because we intended to show that *despite* these multiple, uncorrected comparisons, we still did not find evidence for reliable effects. Overall, our findings challenge the notion that dyslexic and typical readers show differences in FA that are sufficiently systematic to be ascribed to dyslexia (Phase 1), and that these differences reflect various degrees of reading performance (Phase 2).

Interestingly, Vandermosten et al. (2012) identified a significant left temporoparietal cluster in their meta-analysis, which included nine DTI studies using VBA to identify locations where reading ability correlated with FA. Phase 2 of this meta-analysis employed similar methods, yet results are ambiguous. It is worth pointing out that Vandermosten et al. (2012) used GingerALE 2.0.4, which has since been updated to correct major flaws affecting analyses—initial errors made ALE analyses too lenient, therefore inadequately controlling for spurious findings (Turkeltaub et al., 2012). In this study, we used the latest update, GingerALE 2.3.6. The importance of adequate correction for multiple comparisons is striking when visually examining Figures 3 and 4 – virtually all white matter tracts are associated either with comparisons between dyslexics and controls, or as a function of reading ability. This illustrates the heterogeneity of results in the literature, which undoubtedly underlies the absence of robust effect after adequate corrections. More generally, the methods implemented in GingerALE are bound to underestimate inconsistencies between studies (Eickhoff et al., 2009), which indicates that caution should be used when interpreting positive results. In any case, this provides additional strength to the results we report herein – our overall analyses yielded null results *despite* potential biases toward positive results.

There were other differences in statistical analyses between the present meta-analysis and the study by Vandermosten et al. (2012)—these authors did not analyze studies reporting significant differences in FA between dyslexic and typical readers; rather, coordinates of significant differences were integrated with coordinates of significant correlations between reading and FA. Although the intent was to increases statistical power, combining disparate studies can considerably blur the interpretation of the findings. This reluctance to aggregate studies that are fundamentally different was our rationale for distinct analyses. Another point of difference with the Vandermosten et al. (2012) meta-analysis was in our inclusion of two additional studies published after 2012, which in total contained three groups of significant coordinates. This last point in itself does not explain the failed replication, however, given that leaving these studies out still did not allow us to exactly reproduce the findings by Vandermosten et al. (2012).

Our findings are also consistent with a recent study by Koerte et al. (2016), which failed to detect significant differences in FA between dyslexic and typical readers, after correcting for multiple comparisons. The authors proposed that the strict multiple comparison correction employed in their analyses contributed to null results; however, the procedure they used is the recommended one given the high rate of false positives when failing to correct for multiple comparisons (Laird et al., 2005; Moreau, Kirk, & Waldie, 2016; Richlan, 2014; Richlan et al., 2009; Turkeltaub et al., 2012). Together with the present findings, this suggests that differences in other VBA studies may be based on unique characteristics of each individual sample (e.g., demographics, study designs), or false positives, and does not support the idea of systematic differences in FA between dyslexic and typical readers. At the very least, the present findings call for caution when interpreting significant voxels or clusters in a given comparison or correlational analysis, and emphasize the need for statistically sound procedures in future work.

There are several limitations to the present findings. First, we focused on VBA studies, given the consequent body of literature that has shown differences between dyslexics and typical readers based on this method. However, it is important to note that recent studies have used ROI-based approaches, and that these were not included in our analyses. ROI-based approaches allow analyses that can be more theory-driven than whole-brain analyses, yet they are still prone to many issues observed in the VBA literature on dyslexia. For example, the inherent noise of both the data acquisition process and the DTI model (e.g., Zhao, Thiebaut de Schotten, Altarelli, Dubois, & Ramus, 2016), together with flexible pipelines of analysis (Carp, 2012) and failures to correct adequately for multiple comparisons, are all components of the developmental dyslexia literature that potentially exacerbate differences, or could even be responsible for a multitude of spurious findings (see for a recent review Ramus, Altarelli, Jednoróg, Zhao, & Scotto di Covella, 2018). Our focus on VBA studies was deliberate, both for internal consistency and because this type of analysis, applied to developmental dyslexia, has yielded a large number of findings with very little published negative results (Ramus et al., 2018). However, we do not exclude the possibility that more fine-grained methods and analyses could further document structural differences between dyslexics and typical readers (Zhao et al., 2016), although more precise and valid measurements have also been shown to provide additional evidence for a lack of structural differences between groups (Vanderauwera, Vandermosten, Dell’Acqua, Wouters, & Ghesquière, 2015).

In addition, we analyzed together studies that included children and those focusing on adult populations in both phases of this meta-analysis. This was meant to allow generalizations to a disorder, dyslexia, rather than to the same disorder *given specific developmental characteristics*. Averaging these populations together, however, may have reduced the sensitivity of our analyses. For example, age-related reading differences could influence outcomes in children, whereas cortical differences in adult populations could reflect the poorer reading exposure, both quantitative and qualitative, commonly experienced by dyslexic readers throughout their lives (Ben-Shachar, Dougherty, & Wandell, 2007). Aware of this potential limitation, we ran additional analyses separating adults and children to verify that age-related factors did not influence results (see online material for details). In Phase 1, no significant clusters were identified in these separate analyses of children or adults, corroborating initial conclusions. In Phase 2, no significant clusters were found upon analysis of adult studies; however, analysis of positive correlations between reading and FA in five studies including children identified one significant cluster located at MNI coordinates (-18, –8, 34). This suggests a possible positive correlation between reading, as measured by performance on reading-related tasks, and white matter integrity, as measured by FA. It is difficult to confidently relate this finding with dyslexia *per se*, however, given that in three of these studies, participants included a range of typical readers with diverse reading abilities, and the “poor readers” were not explicitly diagnosed with dyslexia (Beaulieu et al., 2005; Nagy, Westerberg & Klingberg, 2004; and Keller & Just, 2009).

It is also possible that specific characteristics of the selected studies disproportionately influenced the results. For example, Zhang et al. (2014) reported coordinates where significant correlations between FA and performance on a Chinese Character Reading Efficiency Test were found among a Chinese readers, yet research has demonstrated that when reading Chinese, a character-based language system, different cortical processing are employed compared to reading English, an alphabetic language system (Siok, Niu, Jin, Perfetti, & Tan, 2008). To ensure that our conclusions were not excessively influenced by the study by Zhang et al. (2014), we re-ran our analysis excluding it. No significant differences were detected in this additional analysis, substantiating our initial conclusions. On a similar note, Nagy et al. (2004) employed unstandardized tests to quantify participants’ reading ability. These may provide less reliable measures of reading ability compared to standardized tests used in other studies included in this meta-analysis. Excluding this study in an additional analysis to determine if results were affected was not possible as the significant coordinate found by Nagy et al. (2004) made up one of two foci in the meta-analysis of negative correlations. Thus, the small number of foci and unstandardized testing may have limited the validity of our results for the negative correlation analysis.

Finally, the number of studies included in our final sample, both in Phase 1 and Phase 2, was rather small. Although this is a legitimate concern, we deliberately favored consistency in the methods reported across studies over sample size, given that small, precise meta-analyses are typically more informative (see for example Turner, Bird, Higgins, Lathlean, & Babidge, 2013). We should also point out that the overall search initially generated a large body of 222 studies, and that Phase 1 included a final sample of 99 and 52 participants, respectively for the two contrasts, whereas Phase 2 included a final sample of 500 and 40 participants, respectively for each correlation analysis. The report of uncorrected clusters for both phases in Figures 3 and 4, as well as the online publication of details for all analyses, are additional attempts to make transparent the process and materials of the present study. Although VBA findings to date do not seem to support fundamental differences in FA associated with reading disability, the addition of future studies might, and these can be easily incorporated into the analyses we reported. Gathering scientific evidence is a cumulative process, and by no means do we imply that the picture we present is definitive, nor do we make specific predictions about future studies. Rather, the present study highlights the lack of current evidence for consistent differences in FA as a function of reading ability, including extreme cases of reading disability.

## 4. Methods and Materials

This section includes two related but distinct meta-analytic searches. The first search (Phase 1) corresponds to the initial contrast between dyslexics and typical readers. It was meant to provide an assessment of FA differences between these two groups. The second search (Phase 2) was intended to explore the relationship between reading ability and FA, within samples of typical readers. The Preferred Reporting Items for Systematic Reviews and Meta-analyses (PRISMA) protocol guided the process and reporting of this meta-analysis in both phases (Moher, Liberati, Tetzlaff, Altman, & The PRISMA Group, 2009). Data, including details about all papers reviewed, spatial coordinates, and all scripts for the reported and additional analyses are freely available online at https://github.com/davidmoreau/2018_Brain_Research

### 4.1 Phase 1

#### 4.1.1 Search

Six databases—PubMed, PsycINFO, ScienceDirect, Scopus, ProQuest Dissertations & Theses Global, and Google Scholar—were searched from 17^th^-21^st^ November 2016 for articles containing relevant material (see criteria below). The search on Google Scholar and ProQuest Dissertation and Theses Global database—which include unpublished dissertations and theses— were intended to minimize publication bias. As all relevant articles were accessible, no additional contact with the authors was required. We used two main search phrases. Firstly, we used the combined search phrases “diffusion tensor imaging and “dyslexia” in PubMed and PsycINFO databases. These search phrases generated too many results when inputted into the remaining databases, ScienceDirect, Scopus, ProQuest Dissertations & Theses Global, and Google Scholar. Therefore, the search phrase was limited to only the title, abstract and keywords for the remaining database. Given this search limitation, synonyms for dyslexia were included. The search phrases used were: “diffusion tensor imaging”, combined with each of the following: “dyslexia”, “reading ability”, “reading disability”, “reading difficulty”, and “reading impairment”.

#### 4.1.2 Eligibility

Several eligibility criteria were employed to eliminate irrelevant articles from database searches when initially scanning the abstract, and later, the full text of articles (see flow diagram, Figure 1). Exclusion criteria included: duplicates, articles in a foreign language, review articles, absence of a comparison between dyslexics and typical readers, studies focusing on reading abilities in typical samples, studies focusing on non-developmental dyslexia, clinical comorbidities, absence of structural comparison, ROI analyses, and methods articles. We detail these criteria hereafter.

Characteristics of participants included in the studies were thoroughly assessed. Given the aforementioned aim of this meta-analysis, studies that did not include a comparison group of dyslexic individuals were excluded. Similarly, we excluded studies in which dyslexia was not explicitly mentioned (e.g., poor reading ability), or those in which participants were not formally diagnosed with dyslexia (e.g., high risk factors but not diagnosis). Furthermore, individuals with pure neglect dyslexia or with spelling impairments only were excluded, as white matter correlates may differ from those of individuals with developmental dyslexia. Studies where dyslexic participants experienced clinical comorbidities, such as TBI, leukemia, pre-term birth or brain tumors, were also excluded because these may confound or provide alternative explanations for any relationships between FA and dyslexia.

Next, studies for which no structural DTI comparisons between dyslexic and control participants, or only including ROI DTI analyses, were excluded. For example, many studies only reported the correlations of FA differences to behavioral measures, such as performance on reading tasks. Methodological studies, such as scanning protocols for children and mathematical modeling, were also excluded. Finally, a handful of studies were excluded because they failed to report spatial coordinates. Details of this procedure are presented in Figure 1.

Overall, 222 articles were identified in the six database searches. Ninety-four were duplicates and were thus removed. The remaining 128 articles were screened based on the abstract, with 89 excluded using the eligibility criteria. The full-text of the remaining 39 articles was assessed for relevance. Thirty-four studies were excluded at this stage, and the remaining five studies were included in the meta-analysis (see Figure 1). More information about studies inclusion/exclusion can be found in the online material.

#### 4.1.3 Data Collection and Analyses

We extracted participant characteristics for each study. This included sample size, gender ratio, groups, mean age and age range, and any other relevant characteristics, such as native language. Then, we logged the coordinates of significant group differences in FA, with the corresponding reference space. The intended summary measures to quantify the outcomes of primary interest were cortical coordinates of voxels/clusters where FA significantly differed between dyslexic and typical reading individuals.

In the studies we selected, coordinates where FA significantly differed were reported in either MNI or Talairach space. Because we conducted the meta-analysis in MNI space, we transformed studies that reported coordinates in Talairach space (2) to MNI space, with the Lancaster et al. (2007) transformation integrated in GingerALE 2.3.6 (Eickhoff et al., 2009, 2012; Turkeltaub et al., 2012). We used GingerALE 2.3.6 software (Eickhoff et al., 2009, 2012; Turkeltaub et al., 2012), which allows ALE implementation. ALE is the most common statistical technique for coordinate-based meta-analyses; the technique converts activation foci into probability distributions centered at specific coordinates. Activation foci are successively centered following a Gaussian probability distribution to generate single modeled activation maps for each study. These maps are then combined in a random-effects model, which provides flexible modeling of uncertainty across studies. Details of the ALE procedure in GingerALE can be found in the online documentation (http://brainmap.org/ale/manual.pdf) and in additional studies (Eickhoff et al., 2009, 2012; Lancaster et al., 2007; Turkeltaub et al., 2012).

Analyses were run separately for coordinates where controls had significantly greater FA than dyslexics (47 foci across 5 experiments), and where dyslexics had significantly greater FA than controls (17 foci across 2 experiments). In both analyses, a false discovery rate (FDR) threshold of .05 was applied (Laird et al., 2005). In addition, given that previous literature has suggested that the reduced and poorer quality reading exposure dyslexic readers likely experience throughout their life may contribute to apparent cortical differences between dyslexic and typical adult readers (Ben-Shachar et al., 2007), we ran separate additional analyses for children and adults. In analyses including coordinates where controls had greater FA than dyslexic readers, three studies included adults and two studies included children. One study including adults and one study including children reported coordinates where dyslexic readers had greater FA than controls.

### 4.2 Phase 2

Phase 1 included studies that specifically explored differences between dyslexic and typical readers. However, it is plausible that differences in FA are subtler and cannot be detected by simply contrasting groups, but could appear when considering a continuum to represent reading ability. With this rationale, many studies have investigated the association between reading ability and FA, regardless of a particular diagnosis of reading disability or dyslexia. To further investigate this relationship, the initial meta-analysis was complemented by a second search, detailed hereafter.

#### 4.2.1 Search

The same six databases used in Phase 1 were accessed and searched during Phase 2 from 24^th^-26^th^ January 2017. In Phase 2 of the study, the initial eligibility criteria were broadened to allow the inclusion of studies comparing dyslexic readers to typical readers via continuous measures spectrum, as well as studies exploring levels of reading ability among non-impaired readers. Eligibility criteria were identical to those of Phase 1, except for the exclusion of studies that did not include dyslexic participants, relevant in the previous search but not in the follow-up. In addition, studies that did not correlate FA with appropriate reading measures were excluded.

In Phase 2, alterations were made to the Phase 1 database searches to reflect the changes in the Phase 2 eligibility criteria. The Phase 1 searches in PubMed and PsycINFO used the narrow keyword of “dyslexia” only, which is unlikely to have generated articles with a range of reading abilities that did not include dyslexia comparisons; thus, these searches were insufficient to generate all required studies for Phase 2. Consequently, the searches in these databases were run again using the broader search phrase “diffusion tensor imaging” combined with each of the following: “dyslexia”, “reading ability”, “reading disability”, “reading difficulty”, and “reading impairment”.

#### 4.2.2 Eligibility

Eligibility criteria were identical to those used in Phase 1, except for the substitution of group comparisons with FA-reading correlations. Firstly, when repeating the searches in PubMed and PsycINFO, the same process as Phase 1 of this study was undertaken. Secondly, studies from Phase 1 searches in ScienceDirect, Scopus, ProQuest Dissertations & Theses Global and Google Scholar that were eliminated because they did not include a dyslexic group, had no structural comparison, or were exclusively ROI analyses were reviewed for inclusion in Phase 2, for a total of 70 studies.

Overall, 171 articles were identified in the six database searches and reviews. Ninety-eight were duplicates and therefore excluded. The remaining 73 articles were screened by reading the abstract and full text where required. A further 64 were excluded based on the exclusion criteria. The remaining nine articles (ten studies in total) were included in the meta-analysis (see Figure 2). More information about studies inclusion/exclusion can be found in the online material.

#### 4.2.3 Data Collection and Analyses

The same participant characteristics and reference space information as in Phase 1 were extracted from the articles. Details of reading task used to measure reading performance, and coordinates where this significantly correlated with FA, were also recorded. The intended summary effect measure to quantify the outcomes of primary interest were cortical coordinates of voxels/clusters where FA significantly correlated with reading ability, as measured by performance on reading tasks.

Coordinates of three studies required transformation from Talairach to MNI space using the Lancaster et al. (2007) transformation in GingerALE 2.3.6 (Eickhoff et al., 2009, 2012; Turkeltaub et al., 2012). Four studies reported findings of significant correlations between FA and multiple reading measures. When coordinates were not directly available, we performed a meta-analysis within individual studies to identify coordinates where FA significantly correlated with overall reading ability, as measured by all reading tasks in the particular study (Eickhoff et al., 2009, 2012; Turkeltaub et al., 2012). Analyses were run separately for coordinates where FA positively correlated with reading ability, and negative correlations. A FDR threshold of .05 was applied, given positive dependence of foci and common conventions (Laird et al., 2005).

Overall meta-analyses were conducted using GingerALE 2.3.6 software (Eickhoff et al., 2009, 2012; Turkeltaub et al., 2012). The overall ALE procedure was identical to this of Phase 1. Analyses were run separately for coordinates where FA positively correlated with reading ability, 42 foci across 9 experiments; and where FA negatively correlated with reading ability, 2 foci across 2 experiments. In both analyses, a FDR threshold of .05 was applied, given independence of foci and typical conventions (Laird et al., 2005). Finally, the meta-analysis was repeated separately for children and adults. Five of the ten studies analyzed included adult participants, whereas the remaining five studies included children. Two studies also reported negative correlations, both of which included child participants. To safeguard against unwarranted generalizations, we ran an additional analysis excluding the study of Zhang et al. (2014), which involved participants reading Chinese text, and compared the findings with those of the overall analysis. All the analyses input and output files can be found in the online material.

